# Cystic fibrosis systemic immune profile is associated with lung microbes and characterized by widespread alterations in the innate and adaptive immune compartments

**DOI:** 10.1101/2023.08.23.553085

**Authors:** Elio Rossi, Mads Lausen, Nina Friesgård Øbro, Antonella Colque, Bibi Uhre Nielsen, Rikke Møller, Camilla de Gier, Annemette Hald, Marianne Skov, Tacjana Pressler, Søren Molin, Sisse Rye Ostrowski, Hanne Vibeke Marquart, Helle Krogh Johansen

## Abstract

Polymicrobial airway infections and detrimental inflammation characterize patients with cystic fibrosis (CF), a disease with heterogeneous clinical outcomes. How the overall immune response is affected in CF, its relationships with the lung microbiome, and the source of clinical heterogeneity are unclear. Our work identifies a specific CF immune profile characterized by widespread hyperactivation, enrichment of CD35^+^/CD49d^+^ neutrophils, and reduction in dendritic cells. Further, our data indicate signs of immune dysregulation due to alterations in Tregs homeostasis, which, together with an impaired B-cell immune function, are linked with patients’ lung function and are potentially the source of clinical heterogeneity. Indeed, clinical heterogeneity does not stem from a specific lung microbiome; yet, commensal bacteria correlate with higher concentrations of circulating immune cells and lower expression of leukocyte activation markers, a condition reversed by pathogenic microorganisms. Overall, our findings provide unique markers and immunomodulatory targets for improving the treatment of CF.

## Introduction

Cystic fibrosis (CF) is a monogenic disease caused by mutations in the *CFTR* gene and is one of the most common autosomal recessive genetic disorders. *CFTR* encodes a chloride channel highly expressed in epithelial cells in various tissues, making CF a multi-organ disease.

CF lung disease is complex and with diverse clinical outcomes^1^. It is characterized by an early onset of chronic inflammation and recurrent bacterial airway infections which is the primary cause of morbidity and mortality. Ongoing pulmonary inflammation and infections lead to structural lung damage and a progressive loss of lung function. It remains unclear whether persistent infections primarily trigger an aberrant inflammatory response or whether it is a precondition that favors microbial colonization. In CF, the immune system fails to resolve the inflammatory response and provide protective immunity against pulmonary infections showing that the CF immune system is highly dysfunctional^2,3^. Thus, flaws in the immune response are directly linked to disease severity in CF.

It is key to understand whether the CF immune system is dysregulated and what mechanisms drive these defects. Innate and adaptive immune cell populations are functionally affected in CF^4^, and these deficiencies have been directly linked to *CFTR* mutations rather than active infections^5,6^. However, the *CFTR* genotype alone cannot explain the heterogeneous disease phenotypes observed in CF patients. Therefore, several other factors, such as mutations in modifier genes, sex, metabolic differences, and the lung microbiota, have been suggested to contribute to the disease phenotype^7^. For example, increased lung microbiome diversity in CF patients correlates positively with lung function^8^. *In vitro*, commensal bacteria isolated from CF airways can reduce the inflammation induced by the common CF pathogen *Pseudomonas aeruginosa* ^9^. Thus, the CF lung microbiome is likely a significant contributing factor modulating the CF immune system, but this remains to be demonstrated.

In this work, using high-resolution flow cytometry and metatranscriptomics analysis on blood and sputum samples, respectively, we investigate the composition and activation status of all immune cell compartments and the transcriptionally active lung microbial communities. Overall, we reconstruct a detailed picture of the immune profile that differentiates CF patients from healthy individuals, potentially identifying alterations in Tregs homeostasis and B-cell responses as the source of the clinical heterogeneity observed in patient’s lung function. Finally, we provide evidence of specific associations between beneficial and pathogenic microbes with a subset of circulating immune cells.

## Results

### Multiple immunological compartments contribute to defining the CF systemic immune profile

To gain insight into the CF immunophenotype, we made a comprehensive whole-blood analysis of immune cell composition and their activation states in CF and healthy subjects (see Methods). Principal component analysis of the absolute concentration (cells/ml) of 82 immune cell subtypes indicates that CF patients cluster separately from healthy subjects (Figure 1A). Diverse branches of the innate and adaptive immune system contribute to the specific CF immune profile, as suggested by the top 10 contributing loadings (Figure 1A). In CF patients, neutrophil concentration is almost twice as high, while dendritic cells (DCs) and CD4^+^ T cell concentrations are 1.8- and 1.2-fold lower, respectively (Figure 1B). All other major immune cell classes show tendencies towards being reduced in CF patients, although not statistically significant.

**Figure 1.**
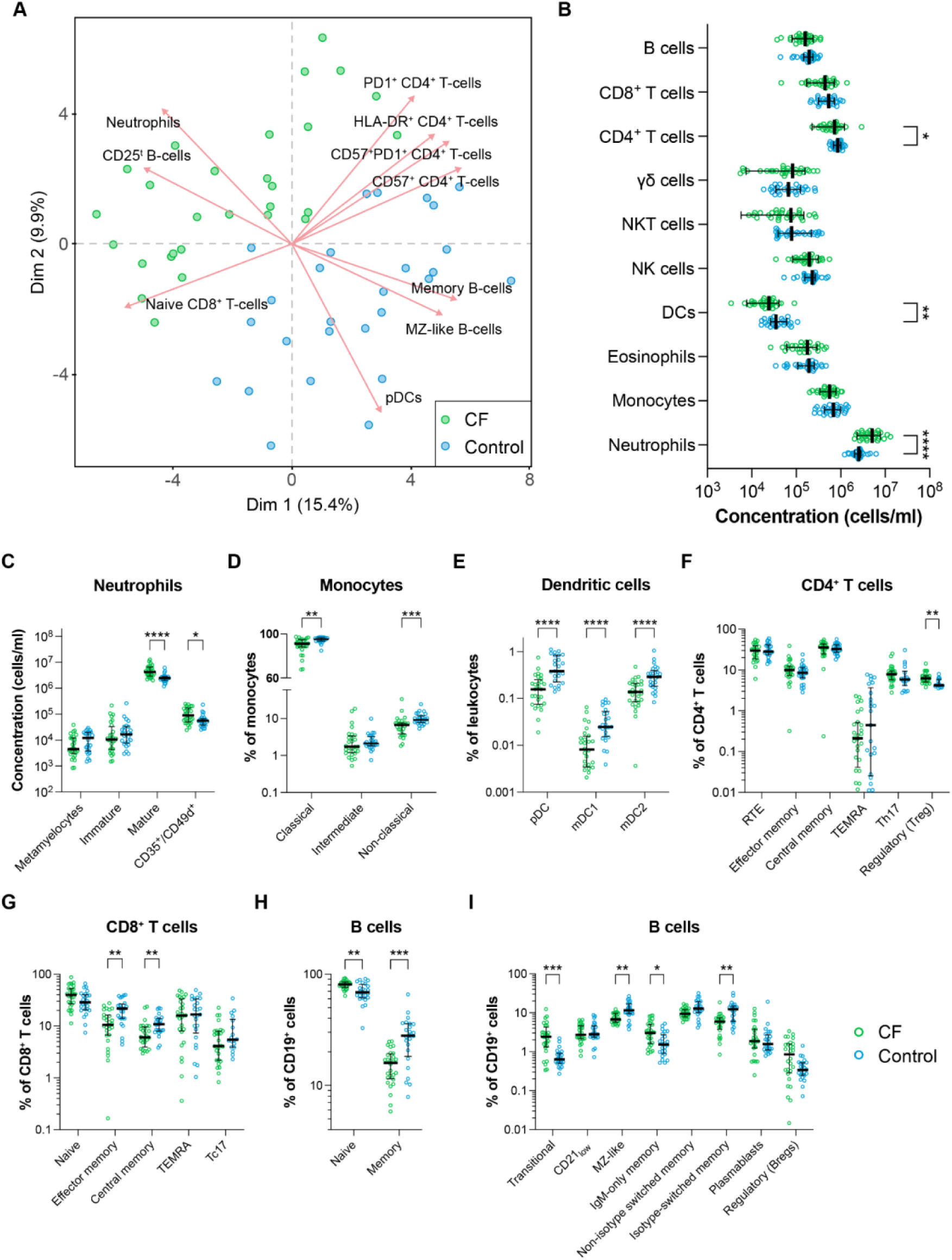
Circulating immune cells characterizing cystic fibrosis patients. **A.** Principal component analysis (PCA) of cell frequencies in whole blood from CF patients and sex- and age-matched healthy controls. The first two principal components (Dim) are plotted. Points represent individual subjects’ data color-coded by subject type (green: CF; blue: control). Loadings of the top 10 contributing variables to Dim1 and Dim2 are shown. **B.** Dot plots represent the median concentration of the major immune cell types (cells/ml). Dots represent the concentration in individual subjects (green: CF; blue: control). Two-sided Wilcoxon test with Benjamini & Hochberg multiple testing correction: *, adjusted *P* value ≤ 0.05; **, adjusted *P* value ≤ 0.01; ****, adjusted *P* value ≤ 0.0001. **C - J**. Dot plots represent the median concentration of specific cell types (cells/ml) or the frequency of parent populations (% of parent cells). Dots represent the value of individual subjects (green: CF; blue: control). Two-sided Wilcoxon test with Benjamini & Hochberg multiple testing correction: *, adjusted *P* value ≤ 0.05; **, adjusted *P* value ≤ 0.01; ***, adjusted P value ≤ 0.001; ****, adjusted *P* value ≤ 0.0001.

Further analysis of the myeloid cell subpopulations shows that the high neutrophil concentration in CF patients is due to increased concentrations of mature CD10^+^ neutrophils and neutrophils with co-expression of CD35^+^/CD49d^+^ (Figure 1C). Further, we observed an overall reduction of both non-classical (CD14^low^CD16^hi^) and classical circulating monocytes (CD14^hi^CD16^low^) in CF patients (Figure 1D). Similarly, plasmacytoid dendritic cells (pDC) and both CD1c^+^ CD141^−^ (mDC1) and CD1c^−^ CD141^+^ (mDC2) myeloid dendritic cells were reduced in CF blood (Figure 1E).

Concerning the acquired cell-mediated immunity, we observed a significant reduction in CD4^+^ T-cell concentration in CF patients (Figure 1B), with a normal distribution of CD4^+^ T-cell subsets except for an increased fraction of FOXP3^+^ regulatory T cells (Tregs) (Figure 1F). In contrast, several CD8^+^ T-cell subpopulations were altered with a decrease in the absolute concentration and fraction of CD8^+^ effector memory T cells, CD8^+^ central memory T cells, and CD161^+^ CD196^+^ CD8^+^ T cells (Tc17 cells), a phenotype associated with IL-17 producing CD8^+^ T cells^10^ (Figure 1G and Suppl. Figure 1C).

Finally, the total B-cell concentration in CF patients was within the normal range of healthy controls (Figure 1B), but CF patients were characterized by a substantially higher fraction of naïve (CD27^−^) B cells paralleled by a reduction in the memory (CD27^+^) B cells (Figure 1H). Further, CF patients had higher fractions of transitional (IgM^+^ CD38^+^ CD27^−^) B cells (Figure 1I), the early, immature B-cell stage that recently emigrated from the bone marrow (Suppl. Table 2). In contrast, the fraction of CD27^+^ marginal zone-like (MZ-like) B cells and isotype-switched memory B cells were reduced in CF patients, whereas the IgM^+^ IgD^−^ memory B-cell subset (IgM-only memory B cells) was elevated (Figure 1I).

### Surface expression of activation markers and check-point molecules indicates hyperactivated immune cells and immune dysregulation

The *CFTR* mutation has been suggested to cause dysfunction of immune cell function and activation^11^. Therefore, we investigated the expression of surface markers involved in the regulation and activation of immune cells in CF patients versus healthy controls.

Several activation markers were upregulated on CF neutrophils. The lectin CD69 (Figure 2A), the complement receptor 1/CD35 (mediating phagocytosis), and the Fc receptor CD64 were increased approximately 1.5 folds on immature and mature neutrophils (Figures 2B and 2C). CD11b, an integrin involved in phagocytosis and adhesion, was more abundantly expressed (1.5-fold) on only immature CF neutrophils (Figure 2B), delineating differences between the two neutrophil subpopulations. The checkpoint molecule Cell Death Protein-Ligand 1 (PD-L1) is induced by proinflammatory stimuli and increased in severe inflammatory conditions such as sepsis^12^. Here, we found the fraction of PD-L1^+^ neutrophils to be twice higher in CF (0.17% vs. 0.075%, *P* value = 0.026) compared to healthy controls (Suppl. Figure 1A). Moreover, the PD-L1 surface expression on PD-L1^+^ neutrophils was 13 times higher in CF patients (MFI: 85.7 vs. 6.6, p < 0.0001) (Figure 2A).

**Figure 2.**
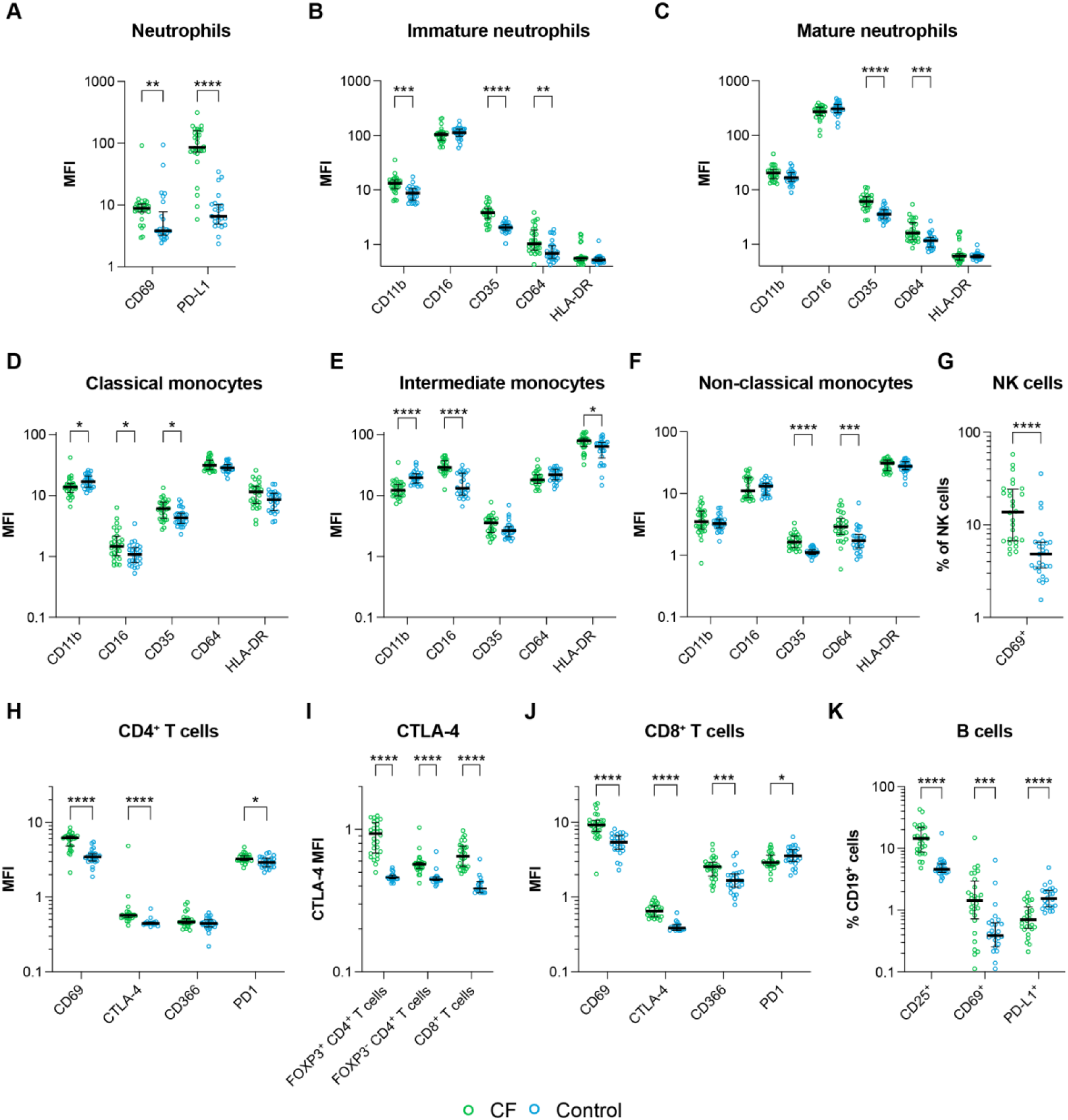
Activation and regulation of immune cells in CF patients and healthy controls. **A – K.** Dot plots represent cell activation or regulatory surface markers’ median fluorescence intensity (MFI) or the median fraction of parent populations (% of parent cells). Dots represent individual subjects (green: CF; blue: control). Two-sided Wilcoxon test with Benjamini & Hochberg multiple testing correction: *, adjusted *P* value ≤ 0.05; **, adjusted *P* value ≤ 0.01; ***, adjusted *P* value ≤ 0.001; ****, adjusted *P* value ≤ 0.0001.

Like neutrophils, circulating CF monocytes displayed an activated phenotype with changes in the expression of several receptors essential for phagocytosis, migration, and antigen presentation. In CF, classical monocytes showed a 1.2-fold decrease in CD11b expression, while expression of CD16 and CD35 receptors increased 1.6 and 1.4 times, respectively (Figure 2E). Similarly, intermediate CF monocytes (CD14^int^CD16^int^) had decreased CD11b (1.7 fold) and increased expression of CD16 (1.9 fold) and HLA-DR (1.3 fold) (Figure 2F). In comparison, non-classical monocytes had increased surface expression of CD35 (1.5 fold) and CD64 (1.7 fold) (Figure 2G). We also observed a 1.4-fold increase of immunomodulator CD366 (TIM-3/T-cell immunoglobulin and mucin-domain containing-3) expression on CF monocytes (Suppl. Figure 1B). The exact role of CD366 in monocytes is not fully understood, but CD366 may be a negative regulator of monocyte IL-12 production with the potential to regulate adaptive Th1 response^13^, which is disfavored in CF patients with chronic *P. aeruginosa* infections^14^, such as those under study (Supplementary Table 1).

Dendritic cells showed no difference, except a decreased expression on mDCs of CD301 (Supplementary Figure 2), a C-type lectin that might be important in regulating exaggerated B-cell responses^15^ and suppression of autoantibodies, which are observed at high frequency in CF patients^16^.

NK cell concentration was normal in CF patients; however, the fraction of activated CD69^+^ NK cells was almost three times higher in CF compared to controls (13.7% vs. 4.8%) (Figure 2H).

Similarly, CF T cells demonstrated an activated phenotype compared to healthy subjects. Non-regulatory FOXP3^−^ CD4^+^ T cells had increased surface expression of the early activation marker CD69 and the immune checkpoint receptors CTLA-4 (cytotoxic T-lymphocyte associated protein-4) and PD1 (Figure 2I). Additionally, we observed a 2.4-fold increase in CTLA-4 levels on regulatory FOXP3^+^ CD4^+^ T cells in CF patients (Tregs) (Figure 2J). While increased expression of CTLA-4 on conventional CD4^+^ T cells is a sign of severe immune activation^17^, high levels of CTLA-4 on Tregs cells underlie the breakdown of Tregs homeostasis, hinting at a state of immune dysregulation in CF patients. Also, CF CD8^+^ T cells showed a difference in surface marker expression with increased expression of CD69 and CTLA-4 together with the checkpoint molecule CD366. However, CD8^+^ T-cell expression of PD1 is reduced in CF (Figure 2K), suggesting that the PD1/PD-L1 axis may be differentially regulated in CD4^+^ and CD8^+^ T cells of CF patients.

Increased immune activation could also be identified in CF B cells. We observed increased fractions of B cells expressing the CD25 and CD69 activation markers (Figure 2L), while the B-cells fraction expressing the PD1 ligand (PD-L1) was reduced in CF patients (Figure 2L). However, the level of PD-L1 expression on the cell surface of PD-L1^+^ B cells was higher in CF (Supplementary Figure 1G), further highlighting the dysregulation in the B-cell compartment and the PD1/PD-L1 immune checkpoint.

### The CF systemic immune profile is linked to lung function

Although a specific peripheral blood immune cell profile characterizes CF patients compared to healthy controls, the clinical phenotypes are often heterogeneous. Thus, we investigated whether the immune profile variations within the CF cohort were associated with patients’ clinical parameters. Average silhouette and gap statistics indicate that CF patients can be split into two groups (C1, n = 15; C2, n = 13) using K-means clustering based on normalized absolute concentration of circulating immune cells (Figure 3A). Comparing CF disease parameters between the two clusters showed that patients in C1 had significantly better lung function (ppFEV_1_) compared to those in C2 (median ppFEV_1_: 73 vs. 41, *P* = 0.0054) (Figure 3B and Supplementary Table 4). No significant differences were observed for other clinical and demographic parameters such as BMI, CF-related diabetes, total IgG levels, *CFTR* mutation, years since the first *P. aeruginosa* isolates, sex, and age (Supplementary Table 4), suggesting a specific link between disease severity heterogeneity and the overall abundance of different immune cell types at a systemic level. C1 is characterized by an increase in the absolute concentration of several immune cell populations belonging to innate and adaptive immunity (Figure 3C). Significant differences could be observed in the B-cell subpopulations, including isotype-switched memory B cells, i.e., cells able to quickly produce efficient pathogen-specific antibodies after antigen reactivation, suggesting an overall increase in cells involved in the protective B-cell response in C1 patients.

**Figure 3.**
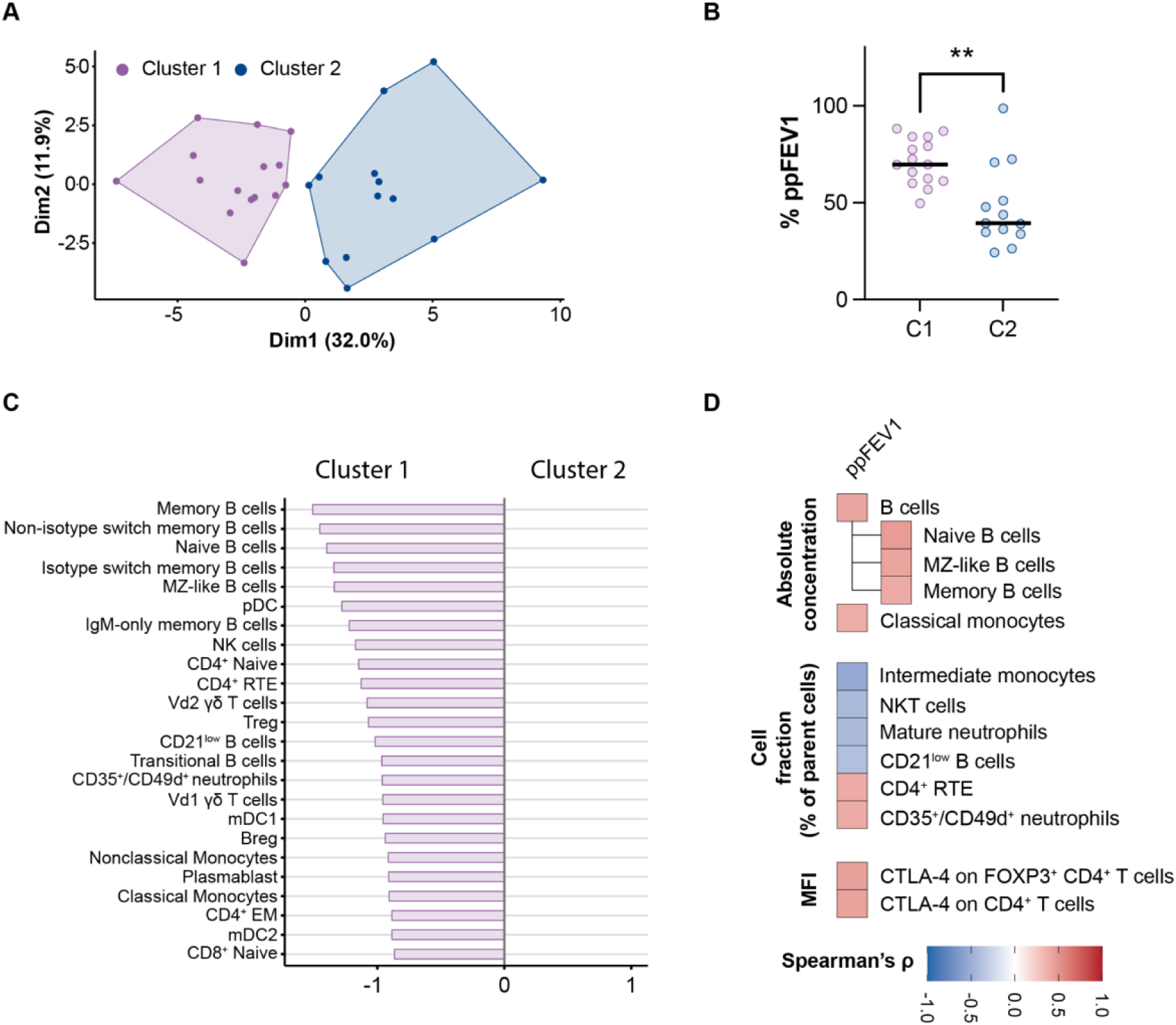
Immunophenotypic variation in the CF cohorts and its relationship with lung function. **A.** CF patient clustering based on K-means clustering using normalized cell concentrations. The optimal number of clusters was identified using average silhouette and gap statistics. **C.** Diverging bar chart representing Z-score differences of significantly different cell populations between patient clusters based on Two-sided Wilcoxon test with Benjamini & Hochberg multiple testing correction. **D.** Correlation heatmaps showing only significant correlations between the lung function predictor ppFEV1 and immune cell absolute concentrations, immune cell frequencies, and surface markers expression. Spearman’s rho rank correlation analysis with *P* < 0.05 was considered statistically significant.

Then, we further explored the dependence of patients’ lung function on the circulating immune cell state, independently correlating the ppFEV1 parameter with the normalized absolute counts and the cell frequencies of all immune cell types, as well as the expression level of all surface markers. As expected, the absolute concentration of B cells, particularly of naïve, MZ-like, and memory B-cells subpopulations and classical monocytes, showed a significant positive correlation with ppFEV_1_ (Figure 3D). Similarly, a higher fraction of the CD35^+^/CD49d^+^ neutrophils subset and CD4^+^ recent thymic emigrants (CD4^+^ RTE) were positively associated with improved lung function (Figure 3D). In contrast, the fraction of natural killer T (NKT) cells, mature neutrophils, intermediate monocytes, and elevated levels of CD21^low^ B cells negatively correlated with ppFEV_1_. Among all activation/regulation markers tested, only the expression of the immune checkpoint regulator CTLA-4 on Tregs and conventional T helper cells correlated positively with ppFEV_1_.

### Systemic immune profiles associate with specific microbes in the lungs but have no direct links with the overall microbiome

The immune system is believed to be influenced by the pathogens actively colonizing CF patients’ airways^18^. Therfore, we sought to explore potential links between the active microbial community in the lungs and the systemic immune system within the CF patients’ cohort. We collected lung expectorates from 26 of the 28 patients and conducted a metatranscriptomics analysis to reconstruct the composition of the transcriptionally active microbiome^19^. The microbiomes recovered were complex, with a median of 55.5 genera (range 28.0 - 62.0) detected in CF expectorates. We observed the co-occurrence of several pathogens commonly associated with CF infections, including bacteria from the *Pseudomonas*, *Stenotrophomonas*, *Mycobacterium*, *Hemophilus*, and *Staphylococcus* genera, as well as fungi from the *Aspergillus* and *Saccharomyces* genera (Supplementary Figure 3 and Supplementary Data 1). At first, we evaluated whether the two groups of CF patients showing different lung functions and immune profiles we identified earlier were characterized by specific microbiomes. PERMANOVA analysis on samples’ β-diversity, calculated using Generalize UniFrac metric, indicated that the transcriptionally active microbiomes did not differ between the two groups (PERMANOVA, *P* = 0.411). Thus, the differences in the overall composition of the local microbial communities have no evident effect in defining the complete immune profiles observed in peripheral blood. However, microbiome composition was significantly associated with ppFEV1 (Mantel test, r = 0.173; *P* = 0.016), indicating a partial contribution of the microbiome to lung function, in agreement with previous observations^20^.

Then, we searched for specific associations between the relative abundance of the most frequent microbial genera (relative abundance of 1% in at least one-fifth of the patients) and different immunological variables, including the absolute concentration of immune cells, distribution of cell subsets, and surface expression levels of activation markers and checkpoint molecules. The relative genera abundances of *Mycobacterium, Staphylococcus*, and the commensal *Veillonella* showed the most significant correlations, whereas the *Pseudomonas* genus showed the lowest (Supplementary Figure 4). Further, based on all significant correlations identified, pathogenic genera (*Mycobacterium*, *Aspergillus, Staphylococcus*) clustered together, except for *Pseudomonas*, which showed correlation patterns more similar to oral commensal genera (*Streptococcus, Veillonella, Prevotella, Rothia*, and *Gemella*) that are known to associate with inflammation negatively^20,21^(Figure 4A-C). Overall, commensal taxa correlated positively with the concentration and fraction of several circulating cells, in particular of those of lymphoid lineage, such as activated CD69^+^ B cells, CD4^+^ and CD8^+^ T-cell subtypes, and regulatory B cells, which were reduced with increasing amounts of the pathogens such as *Staphylococcus, Aspergillus, and Mycobacterium*. Concentration and fraction of PD-L1^+^ B cells, granulocytes, monocytes, CD69^+^ neutrophils, and Tc17 cells positively correlated with the increasing abundance of pathogens while showing a negative correlation with the abundance of commensal genera (Figures 4A and 4B), indicating intra-cohort differences in the PD1/PD-L1 axis.

**Figure 4.**
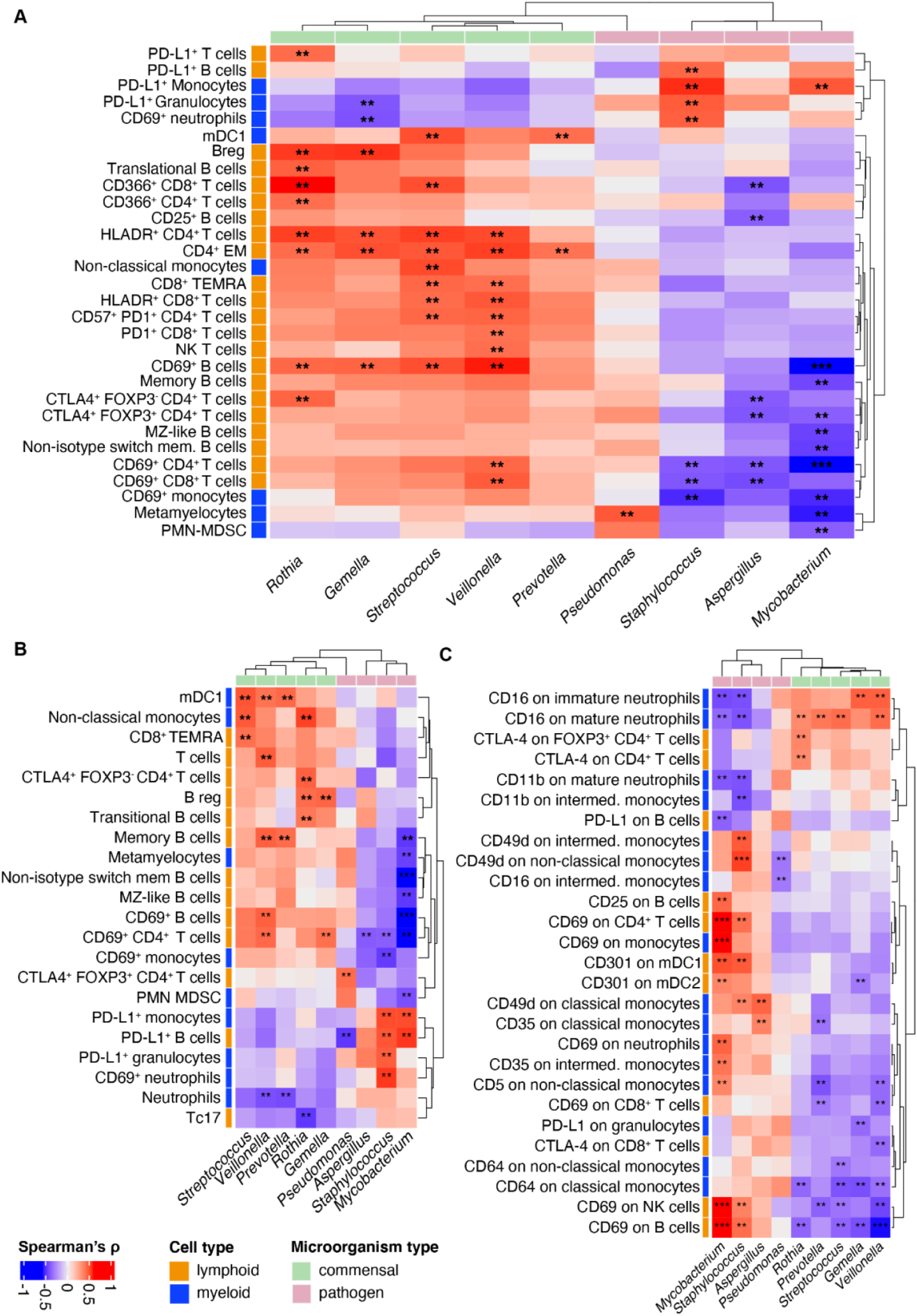
Association between the systemic immune phenotype and the lung microbiome in CF patients. Heatmaps showing all statistically significant correlations (marked with asterisks) between immune cell absolute concentrations (cells/ml, **panel A**), frequencies (% of the parent immune cell type, **panel B**), and median fluorescence intensity (MFI) of cell surface activation markers (**panel C**). Correlations were calculated by Spearman’s rho rank correlation analysis with a *P* value < 0.05 considered statistically significant. *, *P* value < 0.05; **, *P* value < 0.01; ***, *P* value < 0.001. Immune cell type derivation (lymphoid or myeloid) and microorganisms type (commensal or pathogen) are reported.

Similarly, we observed the same tendency with the level of surface markers. Immune activation markers, such as CD69, CD64, CTLA-4, and CD25, and other surface markers, including CD16, CD11b, and CD301, showed an opposite correlation between the abundance of pathogens and commensal bacteria.

## Discussion

In our study, we characterized a specific aberrant CF immune profile and its relationships with the heterogeneity in lung function and the airways microbiome, identifying potential immunological prognostic markers and immune-modulatory targets in both the innate and adaptive immune response.

We defined a specific CF immune profile in peripheral blood with traits common to several autoimmune and chronic inflammatory diseases, which shows widespread immune activation and dysregulation in different immunological compartments suggesting the latter as the source of several clinical manifestations. In our study, we observed several known hallmarks of CF disease, such as increased neutrophil concentrations^22^, a marked reduction in CD4^+^ T cells^23^ (Figure 1B), and a decrease in circulating dendritic cells (DCs), though the latter was previously observed only in mice^24^ and not in humans^25^. Our analysis allowed us to characterize the CF immune profile further. The increase in circulating neutrophils is not limited to mature neutrophils but also depends on a CD49d^+^/CD35^+^ double-positive subpopulation. A proper balance of CD49d^+^ neutrophils is necessary to resolve bacterial infections^26^, while high concentrations of this subpopulation relate to the defective resolution of inflammation^27^ and a predisposition to hypersensitivity reactions^28^, a common condition in CF patients^29^. Although potentially exposing CF patients to co-morbidities, we observed that higher concentrations of CD35^+^/CD49d^+^ co-expressing neutrophils characterize a subgroup of CF patients that showed better lung function and, overall, higher CD35^+^/CD49d^+^ neutrophil frequency positively correlates with ppFEV_1_ (Figure 4).

Immune dysregulation, i.e., the breakdown of the molecular control of immune system processes, leads to disturbed selection or activation of immune cells, impaired regulatory T-cell (Treg) homeostasis, and increased danger signaling in several autoimmune and autoinflammatory diseases. The altered microbiome and defects in the permeability epithelial barrier are emerging factors contributing to immune dysregulation^30^. T-cell responses are known to be dysregulated in CF, with a marked shift toward a Th2- and Th17-dominated response^31^. This effect depends on quantitative and qualitative impairment of Tregs homeostasis occurring after *P. aeruginosa* infections^23^. Although we observed an overall reduction in CD4^+^ T-cells concentrations as previously described, in contrast to previous findings^23^, the Tregs fraction was higher in our CF patients (Figure 1F), and no apparent correlation between the fraction of Tregs and ppFEV1 could be detected, even though most patients (89%) included in our study were defined as chronically infected with *P. aeruginosa*. However, *P. aeruginosa* is the dominant pathogen only in a fraction of patients (Suppl. Figure 2), and we observed that CTLA-4^+^ Tregs absolute concentration and fractions correlated with the relative abundance of different microorganisms, significantly increasing with *Pseudomonas* and decreasing in the presence of *Aspergillus* and *Mycobacterium*. Thus, it can be speculated that the discrepancies observed in the fraction of the Tregs cells might stem from the different active microbial communities at sampling time and suggest that different microorganisms can influence Tregs homeostasis and the delicate balance between adequate pathogen clearance and the development of an uncontrolled inflammatory response^32^.

A central role of Tregs homeostasis in defining CF disease is further supported by the observation that the expression of CTLA-4 is upregulated on Tregs cells in our CF cohort. CTLA-4 is essential for Tregs homeostasis and immuno-suppressive function, thus indicating a specific immune dysregulation that might account for the reduction of a beneficial immune response and increased pathogen persistence^32^. Nevertheless, our data indicated that CTLA-4 expression level, and thus its regulatory function on Tregs and classic CD4^+^ T cells, positively correlate with patients’ ppFEV1 parameter and, thus, with lung function. Interestingly, CTLA-4 upregulation on the two cell types seems to be significantly linked with the relative abundance of the *Rothia* genus. *Rothia mucilaginosa* can reduce inflammation by modulating the NF-kB pathway^33^, which is also involved in regulating Tregs development and function and might be the source for the CTLA-4 changes we observed in our CF cohort.

The CF immune phenotype is also characterized by quantitative and qualitative changes in circulating dendritic cells (DCs). DCs absolute concentration was significantly reduced in CF patients, which resembles other respiratory diseases characterized by chronic inflammation and recurrent microbial infection^34^. Further, the expression of the C-type lectin CD301 on myeloid dendritic (mDC1/mDC2) is lower in CF. Although generally reduced in CF patients, CD301 levels on mDCs might also be modulated in response to active microbes: CD301 expression correlates positively with the relative abundance of pathogenic taxa, such as *Mycobacterium* and *Staphylococcus*. In contrast, the marker expression is significantly reduced with the increasing abundance of commensal taxa, specifically with the *Gemella* genus. Depletion of CD301^+^ DCs relates to an impaired Th2 response upon nematode infections^35^, and CD301^+^ DCs are necessary for IL-17 production from TCRγδ T-cells and Th17 cells following epidermal infection with *Candida albicans* or intranasal infection with *Streptococcus pyogenes*^36,37^. Thus, the overall reduction of CD301 and CD301^+^ DCs might contribute to Th17-skewed production of the proinflammatory cytokine IL-17 and modulation of the Th1 and Th2 responses.

CD301 expression by dendritic cells also plays a critical role in B-cell maturation and activity. CD301 depletion correlates with the increasing generation of autoreactive antibody responses^15^, which are found in up to 80% of CF patients and are linked to worse prognosis^16^. Although we could not verify this correlation in our study cohort, we did observe significant dysfunction of the B-cell compartment in CF patients. Despite having absolute B-cell concentrations comparable to healthy subjects, CF patients showed reduced MZ-like and isotype-switched memory B-cells fractions and increased fractions of naïve B cells and IgM-only memory B cells. These characteristics are also observed in chronic lung diseases, such as bronchiectasis and scleroderma lung disease^38,39^, as well as in individuals affected by common variable immunodeficiency (CVID)^40^, a primary immunodeficiency disease with reduced ability to isotype-switch and recurrent airways infections due to low levels of protective antibodies^40^. CF patients’ reduced capacity for isotype-switch and development of memory may compromise the high-affinity secondary antibody responses, contributing to the increased susceptibility to recurrent opportunistic infections.

Our study identified significant differences in the immune profiles of CF patients, which may offer potential targets for clinical interventions. However, clinical outcomes in CF patients are often heterogeneous, particularly at the level of lung function. B-cell function might be particularly relevant in defining these differences (Figure 3). Indeed, we identified two groups of CF patients showing a remarkable divergence in the ppFEV1 parameter, with those displaying a better lung function having higher absolute concentrations of several B-cell subtypes. Thus, although B cells are similar between healthy controls and CF patients when taken as a whole, within the CF cohort, some patients may maintain a better B-cell response associated with a lower decline in lung function.

However, an increased fraction of the CD21^low^ B-cell population was associated with a worse lung function. The CD21^low^ subset is expanded in conditions characterized by chronic immune stimulation, as frequently observed in CVID patients^41^, where CD21^low^ cells are associated with immune dysregulation and autoimmune disease^42^. Thus, elevated fractions of CD21^low^ B cells may indicate immune dysregulation in CF, representing another player in determining diversity in patients’ clinical outcomes, as described in CVID patients^43^.

Whether the differences we observed in CF patients are intrinsic or acquired due to the infection history cannot be clarified by our work. The role of the microbiome in CF and inflammation is still debated^18^. Commensal bacteria seem to reduce hyperinflammation induced by *P. aeruginosa*^9^. At the same time, enrichment of the lung microbiome with oral taxa is associated with a Th17-dependent inflammatory response^44^.

Our data suggest that changes in the microbial community can play a significant role in disrupting the balance of the immune response. Commensal bacteria and opportunistic pathogens showed distinct and opposite correlation profiles for several immunological variables analyzed: *Prevotella*, *Rothia, Veillonella,* and *Gemella* positively correlated with the abundance and frequency of several lymphoid cells that are also enriched in patients with higher ppFEV1 levels. However, the abundance of commensals negatively correlated with several activation markers on both myeloid and lymphoid cells. On the contrary, the expression of the same markers increased with the relative abundance of pathogenic taxa. Although we could not prove that the correlations we observed were all biologically relevant, in agreement with previous studies^20,21,33^, our data suggest that oral commensals promote the activity of the overall immune system and, at the same time, avoid immune hyperactivation connected with the increasing presence of pathogenic taxa.

Among the pathogenic taxa, the *Pseudomonas* genus, however, represents an interesting case. We observed a positive correlation between *Pseudomonas* and metamyelocytes, but we did not detect any significant correlation with other immunological variables, despite the known role of *P. aeruginosa* in CF pathogenesis. This discrepancy may be attributed to several factors. The bacterium chronically infects substantial fractions of CF patients in our cohort, potentially masking changes in immune variables. Additionally, the impact of *P. aeruginosa* may be minimal at the systemic level. As we solely investigated circulating immune cells, changes at the site of infection may have gone unnoticed.

Despite the efficacy of CFTR modulators in reducing traditional CF pathogens, a significant proportion of patients remain infected^45^. This underscores the pressing need for a more comprehensive understanding of host-pathogen interactions in CF. Our work highlights how the overall immune system is affected in CF, identifying several traits similar to other patients suffering from immunity errors, especially immune dysregulation, and provides a crucial foundation for developing evidence-based medicine in CF management. Further, our observations suggest that the coexistence of various members of microbial communities may influence the immune response in CF. Consequently, any immune modulatory intervention should acknowledge the complex interactions with the microbiome occurring during the treatment.

## Online methods

### Study cohort

Twenty-eight CF patients attending the Copenhagen CF center at Rigshospitalet were consecutively recruited (Supplementary Table 1). CF patients were predominantly males (61%) with a median age of 34 years (range 12 - 61). Most CF patients were characterized by homozygous *CFTR* ΔF508 mutation (67.9%), while only 28.6% had a secondary mutation in addition to ΔF508. All mutations were classified as severe. Almost all patients were clinically defined as chronically infected by *Pseudomonas aeruginosa* (89.3%, Copenhagen criteria^46^) and had a median percent predicted forced expiratory volume in 1 s (ppFEV_1_) of 62% ranging between 24 – 99%.

In addition, 27 age- and sex-matched healthy controls were included anonymously from the Blood Bank, Department of Clinical Immunology at Rigshospitalet (Supplementary Table 1).

### Samples collection and pre-processing

Peripheral blood samples were collected in three separate tubes (K_2_EDTA, 3 ml; Sodium Citrate, 3.5 ml; Lithium Heparin, 4 ml) by venipuncture and processed for flow cytometry staining the same day. At the same time, sputum samples were collected and processed as previously described^19^, with minor modifications. Briefly, expectorates were collected directly from a patient and immediately added to freshly prepared sputum pre-lysis and preservation buffer (3 ml 1× DNA/RNA shield per 1–2 ml sputum sample, Zymo Research, USA; 200 mM Tris(2-carboxyethyl)phosphine, TCEP; 100 μg/ml Proteinase K, Thermo Fisher Scientific, USA) and vigorously shaken by hand until the samples were homogenous and wholly lysed. Samples were briefly stored at 4 °C and then at −80 °C for long-term storage.

### Peripheral blood immune cell phenotyping (flow cytometry)

A highly standardized, customized DuraClone antibody panel^47^ (Beckman Coulter, USA) especially developed for immune cell profiling of lymphoid and myeloid cell subsets, was used. Whole blood was labeled according to the manufacturer’s instructions using seven tubes containing each a lyophilized 10-plex antibody cocktail. A complete list of the antibodies used is provided in Supplementary Table 2. The first tube contained lineage-specific antibodies and acquisition beads to assess absolute cell counts, which were used to calculate absolute concentrations of cell subsets in the remaining tubes. The samples were analyzed on a Navios Ex flow cytometer (Beckman Coulter, USA). The acquired data were analyzed using serial gating strategies in standardized predefined analysis templates in Kaluza software 2.1 (Beckman Coulter, USA). A complete list of gated cell populations and their marker combinations can be found in Supplementary Table 3. Data exported from Kaluza were processed in a custom script to calculate absolute concentrations, after which visualization and statistical analyses were performed using R software (version 4.2.1) and GraphPad Prism (version 9.4.1).

Gated cell populations were analyzed by principal component analysis (PCA) using the R package “FactorMineR”. Missing data were imputed by a PCA method using the R package “missMDA”. Hierarchical cluster analysis and K-means clustering were performed on log_10_-transformed data using “pheatmap” and “stats” package in R, respectively. Hierarchical clustering was performed using the complete agglomeration method with Euclidian distances. Categorical variables were compared using the χ^2^ test. Correlations were calculated using Spearman’s rho rank correlation analysis using the “Hmisc” R package.

### Sputum sample processing and sequencing library preparation

Frozen sputum samples were thawed at room temperature. After adding 1 mg/mL lysozyme, samples were incubated for 10 minutes at room temperature and then for another 10 minutes on ice. Samples were homogenized in ZR BashingBead Lysis Tubes (Zymo Research, USA), performing 3 homogenization cycles (30 seconds at 6,500 rpm and 2 minutes incubation on ice) using Precellys 24 homogenizer (Bertin Instruments, USA). Total RNA was extracted using Quick-DNA/RNA Miniprep (Zymo Research, USA). Between 0.8 – 5 µg of total RNA was digested with 6 – 10 U of TURBO DNase (Thermo Fisher Scientific, USA) for 30 minutes. DNAse-digested RNA was purified using the RNA Clean & Concentrator-5 kit (Zymo Research, USA), selecting RNA species larger than 200 bp. The recovered RNA was quantified using fluorometric quantitation, and the fragmentation state (DV_200_) was evaluated using an RNA Nano kit on an Agilent Bioanalyzer 2100 machine (Agilent Technologies). All samples with a DV_200_ between 50% and 97% were further processed. Between 250 ng and 1 µg of DNAse-digested samples were depleted of rRNA species using a custom combination of riboPOOL Human:Pan-Bacteria (78:22 ratio) kits (siTOOLs Biotech, Germany). rRNA-depleted samples were used to prepare strand-specific sequencing libraries using the KAPA RNA HyperPrep Kit (Roche, Switzerland). After optimization, the fragmentation step time was reduced to 3 min to overcome the partially fragmented nature of the samples. Sequencing was performed on an Illumina NextSeq 500 machine generating a minimum of 150 million reads per sample of either 1 x 75 bp or 2 × 75 bp reads.

### Lung microbiome analysis

Low-quality bases and contaminant adapters were trimmed using Trimmomatic (v 0.35), discarding reads shorter than 35 nt (minimum length to avoid excessive human reads contamination in meta-transcriptomes). Reads were further processed using the SortMeRNA tool (v 2.1) to remove reads generated from residual rRNA transcripts. High-quality human and bacterial reads were separated *in silico* by mapping reads using the BWA aligner and MEM algorithm against the human genome assembly GRCh38.p9 retrieved from the NCBI database. Reads not mapping on the human genome were used as input for analyzing the transcriptionally active bacterial community identifying bacterial genera using the KRAKEN tool (version 2.1.2). Kraken output was processed with the Bracken tool (version 2.6.2) to obtain the final Operation taxonomic units (OUTs) tables. Taxa with less than 5 predicted fragments assigned were removed, and fragments counts were used to calculate each taxon’s fraction in the samples. Taxa less than 0.1% in at least 6% of the samples (n = 3) were discarded. Normalized fractions were used to calculate samples’ beta-diversity based on Generalized UniFrac (Chen *et al.*, 2012) (with alpha 0.5) metric using the GUniFrac() function from GUniFrac R package and a phylogenetic tree generated using ETE Toolkit^48^. Permutational multivariate ANOVA (PERMANOVA) and Mantel test, as implemented in the “vegan” R package, were used to evaluate relationships between respiratory bacterial composition (β-diversity) and patients’ immune groups and ppFEV1 parameter, respectively. Spearman’s rho rank correlation analysis was used to compute correlations between microbial taxa’s relative abundance and immune cells’ abundance and fraction.

## Ethical approval and consent to participate

The local ethics committee approved using the samples at the Capital Region of Denmark Region Hovedstaden (registration number H-19001151, approved 07/03/2019), and all patients gave informed consent according to the current laws.

## Author contributions

ER, SRO, HKJ, and HVM designed the study. HVM designed the immune flow panel and implemented the highly standardized flow setup method. ER, NFØ performed the experiments with the additional help of AC. ER, ML, HVM, and NFØ accessed, verified, and analyzed the data. ML, ER, NFØ, VB, HKJ, SM, HVM, and SRO contributed to data interpretation. BUN, RM, CG, HKJ MS, and TP recruited the patients. ER, RM, BUN, CG, and AH collected patients’ samples and information. ER, ML, NFØ wrote the manuscript with the input of all authors. All authors approved the final manuscript.

## Data Sharing

Raw sequence read data supporting the results of this work are available in the EMBL-EBI European Nucleotide Archive (ENA) under Accession No. PRJEB56242.

## Acknowledgment

This research was funded by The Novo Nordisk Foundation Project Grants in Bioscience and Basic Biomedicine (grant n. NNF18OC0052776) and the Danish Research Council (DFF-9039-00037A) grants awarded to HKJ. ER was supported by a Cariplo Foundation “Biomedical research conducted by young researchers” grant n. 2020-3581.

