## Supplementary Material for "Cystic fibrosis systemic immune profile is associated with lung microbes and characterized by widespread alterations in the innate and adaptive immune compartments"

**by**

Elio Rossi, Mads Lausen, Nina Friesgård Øbro, Antonella Colque, Bibi Uhre Nielsen, Rikke Møller, Camilla de Gier, Annemette Hald, Marianne Skov, Tacjana Pressler, Søren Molin, Sisse Rye Ostrowski, Hanne Vibeke Marquart, Helle Krogh Johansen

Supplementary Table 4…………………………………………………………...16

### SUPPLEMENTARY FIGURES

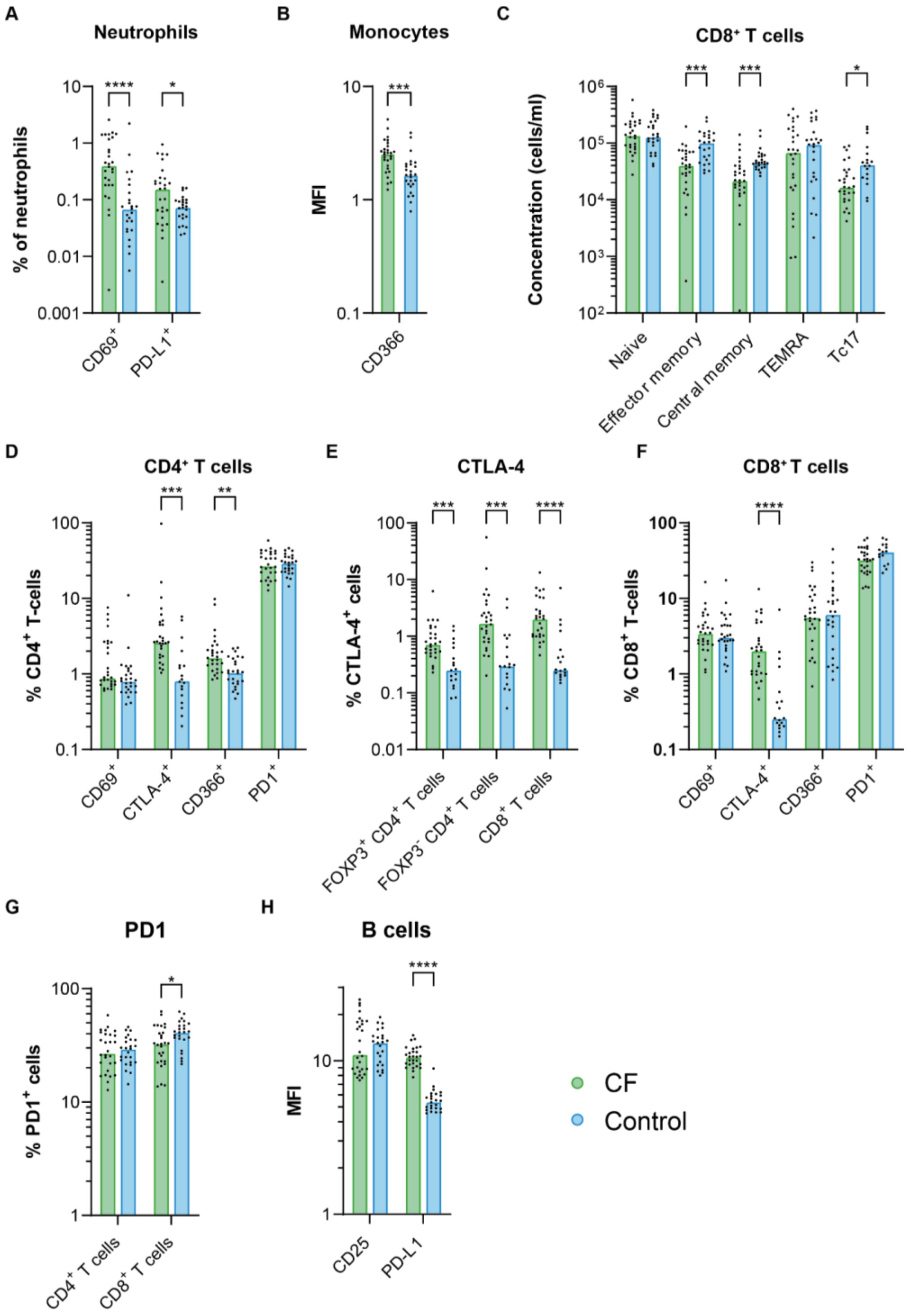

**Supplementary Figure 1. Immunophenotypic differences between cystic fibrosis (CF) patients and healthy controls. A-H.** Barplots represent the median concentration, frequency, or MFI immune variables. Dots represent individual subjects. Two-sided Wilcoxon test with Benjamini & Hochberg multiple testing correction: *, adjusted $P$ value < 0.05; **, adjusted $P$ value < 0.01; ***, adjusted $P$ value < 0.001; ****, adjusted $P$ value < 0.0001. Green: CF, blue: Controls. **A.** Frequency of CD69- and PD-L1 positive neutrophils, **B.** Expression of CD366 on monocytes, **C.** Concentration of $CD8^+$ T-cell subsets, **D.** Frequency of $CD4^+$ T cells expressing activation/inhibitory receptors, **E.** Frequency of CTLA4 expressing T cell, **F.** Frequency of $CD8^+$ T cells expressing activation/inhibitory receptors, **G.** Frequency of T cells expressing PD1, **H.** Expression of CD25 and PD-L1 on B cells.

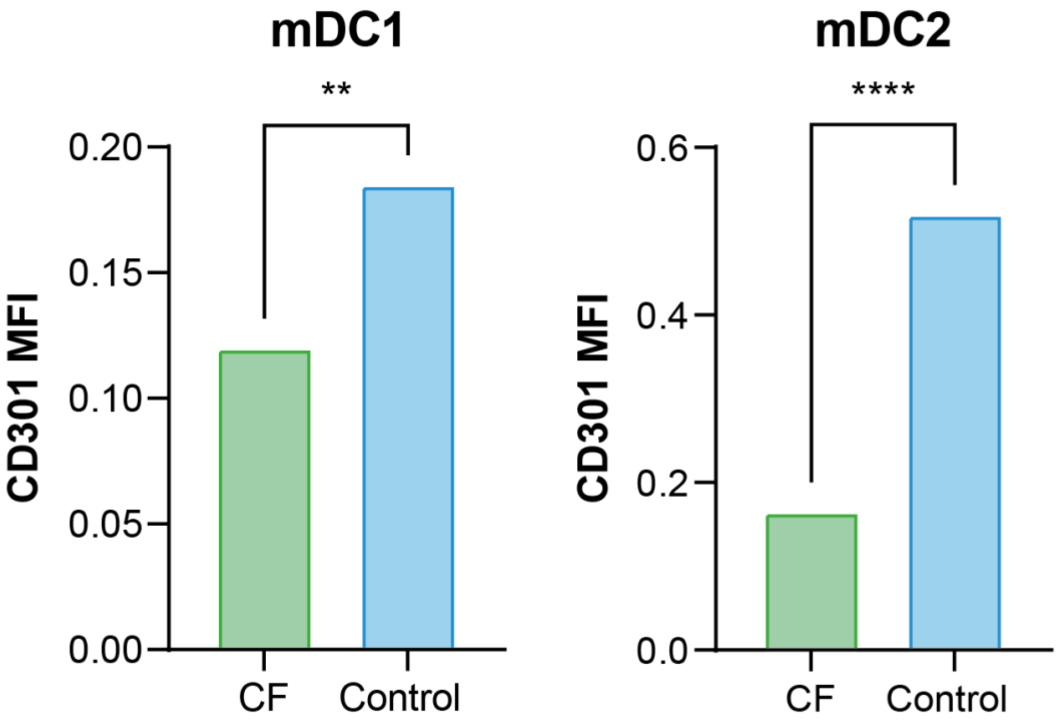

**Supplementary Figure 2. Surface expression of CD301 marker in cystic fibrosis (CF) patients and healthy controls in myeloid. A-H.** Barplots represent the expression (median fluorescence intensity, MFI) of the CD301 marker on $CD1c^{+}$ $CD141^{-}$ (mDC1) and $CD1c^{-}$ $CD141^{+}$ (mDC2) myeloid dendritic cells in cystic fibrosis patients (CF, green) and healthy control (Control, blue). Two-sided Wilcoxon test with Benjamini & Hochberg multiple testing correction: **, adjusted *P* value < 0.01; ****, adjusted *P* value < 0.0001.

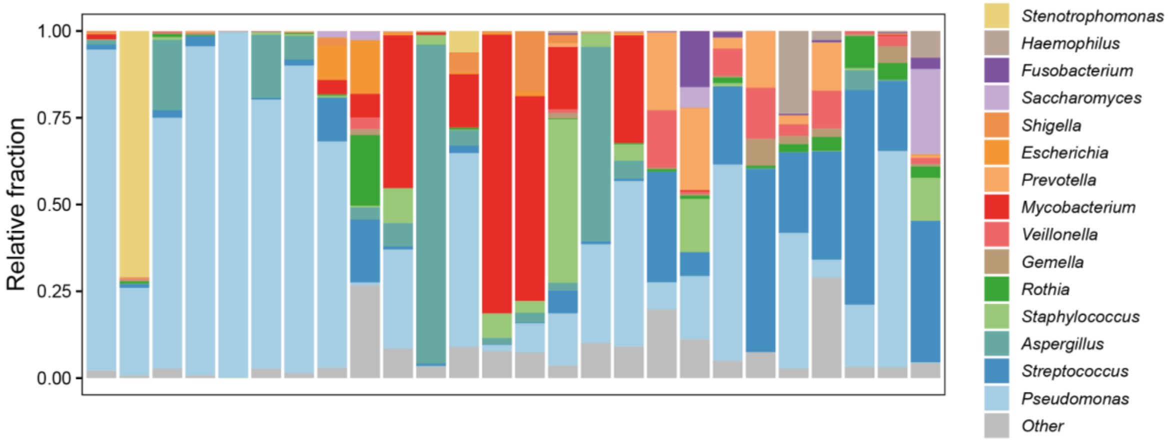

**Supplementary Figure 3. Local lower airway microbiome composition based on microbial transcriptional activity.** Genus-level classification of microorganisms obtained from the analysis of RNA sequences from cystic fibrosis patients' expectorates. Bars represent relative abundances of different genera in samples from individual patients. Only genera with average relative abundances of > 1% in 20% of the patients are shown. The remaining genera are binned in the group "Other". A complete list of all genera identified is reported in Supplementary Data 1.

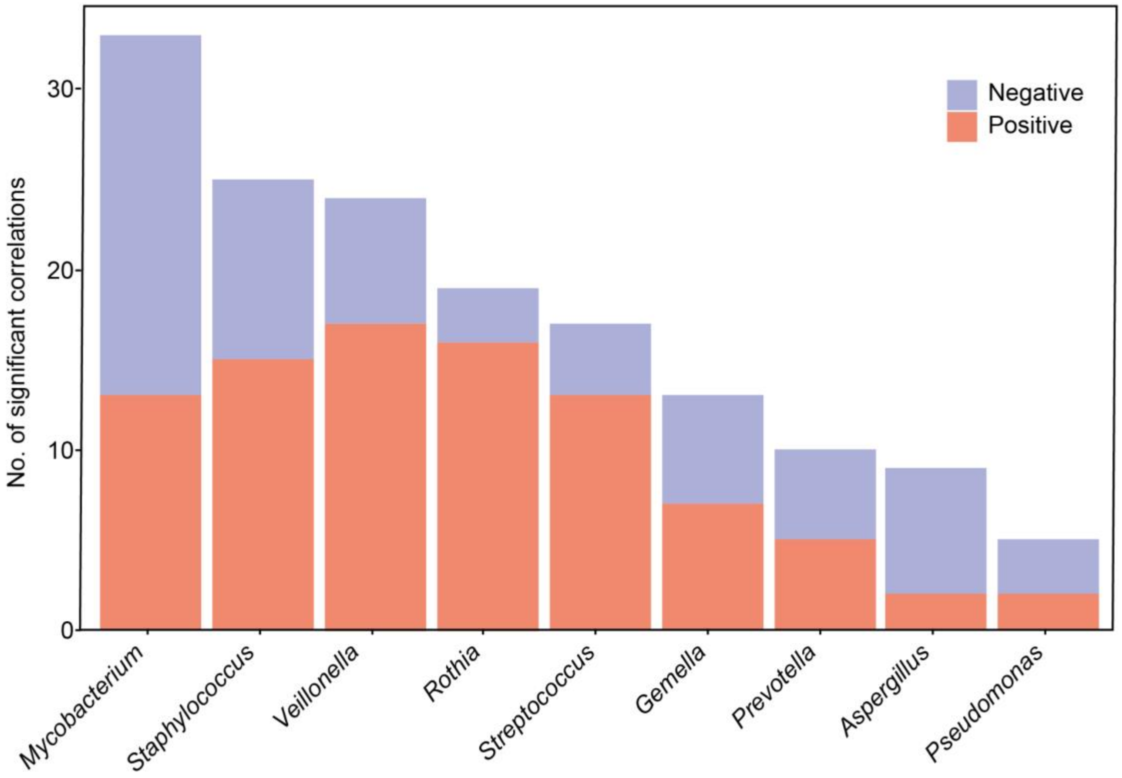

**Supplementary Figure 4. Correlation between the transcriptionally active microbiome and systemic immune variables**. Bars represent the total number of statistically significant correlations between any immunological variable and the relative abundance of each top microbial genus (relative abundance >1% in at least 20% of patients) in cystic fibrosis patients. Correlations were calculated using Spearman's rho rank correlation analysis, and a $P$ value $< 0.05$ was considered statistically significant. The bars are color-coded based on the direction of the correlation (blue, top bars: negative correlation; red, bottom bars: positive correlation).

### SUPPLEMENTARY TABLES

**Supplementary Table 1. Clinical characteristics of cystic fibrosis (CF) patients and control cohort in the study.**

| **Demographics** | **CF cohort** | **Control cohort** | ***P* value** | **Test** |
|---|---|---|---|---|
| All (%) | 28 (100) | 27 (100) | | |
| Sex | | | | |
| Female, n (%) | 11 (39.3) | 15 (55.6) | 0.285 | Fisher’s exact |
| Male, n (%) | 17 (60.7) | 12 (44.4) | | |
| Age, median (range) | 34 (12 - 61) | 36 (24 - 65) | 0.167 | Mann-Whitney |
| FEV1%, median (range) | 62 (24 - 99) | .. | | |
| Mutation | | | | |
| F508 homo, n (%) | 19 (67.9) | .. | | |
| F508 hetero, n (%) | 8 (28.6) | .. | | |
| Other, n (%) | 1 (3.6) | .. | | |
| Chronic *P. aeruginosa*, n (%) | 25 (89.3) | .. | | |

**Supplementary Table 2. Antigens targeted for cell characterization by flow cytometry.**

| Tube | FL1 | FL2 | FL3 | FL4 | FL5 | FL6 | FL7 | FL8 | FL9 | FL10 |
|---|---|---|---|---|---|---|---|---|---|---|
| **Lineage** | CD61 | CD69 | CD16 | CD56 | CD19 | CD8 | CD4 | CD14 | CD45 | CD3 |
| **B cells** | IgG | CD21 | CD27 | CD1d | PD-1L | CD38 | CD5 | IgM | CD19 | CD24 |
| **T cells** | CD45RA | CCR7 | CD28 | PD-1 | CD27 | CD31 | CD4 | CD57 | CD8 | CD3 |
| **TCR subsets** | TCRγδ | TCRαβ | HLA-DR | CD366 | TCRvδ2 | CD8b | CD4 | TCRvδ1 | CD8a | CD3 |
| **Tregs/Th17** | CD127 | CD25 | CTLA4 | CD39 | CD196 | FoxP3 | CD4 | Helios | CD3 | CD161 |
| **Dendritic cells** | CD141 | LIN | CD304 | CD1c | CRLF2 | CD301 | CD11c | HLA-DR | CD16 | CD123 |
| **Myeloid cells** | CD35 | CD49d | CD10 | Siglec-8 | CD64 | CD66b | CD16 | HLA-DR | CD14 | CD11b |

**Supplementary Table 3. Flow cytometric characterization of immune cell populations**

| Name | Marker combination | Frequency (%) | MFI |
|---|---|---|---|
| **Tube 1: CD61/CD69/CD16/CD56/CD19/CD8/CD4/CD3/CD14/CD45** | | | |
| Leukocytes | CD45pos | | |
| Neutrophils | CD16pos CD45pos SShigh | % of CD45pos | |
| Activated neutrophils | CD69MFI CD16pos CD45pos SShigh | % of neutrophils | CD69_MFI |
| Eosinophils | CD16neg CD45pos SShigh | % of CD45pos | |
| Monocytes | CD14pos CD45pos | % of CD45pos | |
| Activated monocytes | CD69MFI CD14pos CD45pos | % of monocytes | CD69_MFI |
| Lymphocytes | CD45pos Sslow | % of CD45pos | |
| T-cells | CD3pos | % of lymphocytes | |
| CD4+ T-cells | CD4pos CD3pos | % of lymphocytes | |
| Activated CD4+ T-cells | CD69MFI CD4pos | % of CD4+ T-cells | CD69_MFI |
| CD8+ T-cells | CD8pos CD3pos | % of lymphocytes | |
| Activated CD8+ T-cells | CD69MFI CD8pos | % of CD8+ T-cells | CD69_MFI |
| Mature B-cells | CD45bri CD19pos | % of lymphocytes | |
| Activated mature B-cells | CD69MFI CD45bri CD19pos | % of mature B-cells | CD69_MFI |
| NK-cells | (CD16pos and CD56pos) CD3neg | % of lymphocytes | |
| Activated NK-cells | CD69MFI (CD16pos and CD56pos) | % of NK-cells | CD69_MFI |
| NKT-cells | CD56pos CD3pos | % of lymphocytes | |
| **Tube 2: IgD/CD21/CD27/CD274/CD38/CD19/IgM/CD24/CD25/CD5** | | | |
| B-cells | CD19pos | % of lymphocytes | |
| Mature naive B-cells | IgDpos IgMpos CD27neg CD19pos | % of CD19pos | |
| Memory B-cells | CD27pos CD19pos | % of CD19pos | |
| Isotype-switched B-cells | IgDneg IgMneg CD19pos | % of CD19pos | |
| Isotype-switched memory B-cells | IgDneg IgMneg CD27pos CD38dim CD19pos | % of CD19pos | |
| Transitional B-cells | IgMbri CD38bri CD27neg CD19pos | % of CD19pos | |
| IgM only memory B-cells | IgDneg IgMpos CD27pos CD19pos | % of CD19pos | |
| Marginal Zone like B-cells | IgDpos IgMneg CD27pos CD19pos | % of CD19pos | |
| Plasmablasts | CD27bri CD38bri CD19pos | % of CD19pos | |
| Bregs | CD24bri CD38bri CD19pos | % of CD19pos | |
| CD21low B-cells | CD21low CD38dim CD19pos | % of CD19pos | |
| Activated B-cells | CD25MFI CD19pos | % of CD19pos | CD25_MFI |
| PD-L1pos B-cells | CD274MFI CD19pos | % of CD19pos | PD-L1_MFI |
| Granulocytes | SSh | | |
| PD-L1pos Granulocytes | CD274MFI SSh | % of granulocytes | PD-L1_MFI |
| Monocytes | FSm SSm | | |
| PD-L1pos Monocytes | CD274MFI FSm SSm | % of monocytes | PD-L1_MFI |
| **Tube 3: CD45RA/CD197/CD27/CD279-PD1/CD28/CD31/CD4/CD3/CD57/CD8** | | | |

| Name | Marker combination | Frequency (%) | MFI |
|---|---|---|---|
| T-cells | CD3pos | % of lymphocytes | |
| CD4 T-cells | CD4pos CD3pos | % of lymphocytes | |
| Recent Thymic Emigrants (RTE) CD4 T-cells | CD45RApos CD31pos CD4pos | % of CD4pos | |
| Naive CD4 T-cells | CD45RApos CD197pos CD4pos | % of CD4pos | |
| Central memory CD4 T-cells | CD45RAneg CD197pos CD4pos | % of CD4pos | |
| T-effector memory RA (TEMRA) CD4 T-cells | CD45RApos CD197neg CD27neg CD28neg CD4pos | % of CD4pos | |
| Effector memory CD4 T-cells | CD45RAneg CD197neg CD4pos | % of CD4pos | |
| Terminal differentiated CD4 T-cells | CD57pos CD4pos | % of CD4pos | |
| Exhausted terminal differentiated CD4 T-cells | CD57pos CD279pos CD4pos | % of CD4pos | |
| PD1pos CD4 T-cells | CD279MFI CD4pos | % of CD4pos | PD1_MFI |
| CD8 T-cells | CD8pos CD3pos | % of lymphocytes | |
| Naive CD8 T-cells | CD45RApos CD197pos CD8pos | % of CD8pos | |
| Central memory CD8 T-cells | CD45RAneg CD197pos CD8pos | % of CD8pos | |
| T-effector memory RA (TEMRA) CD8 T-cells | CD45RApos CD197neg CD27neg CD28neg CD8pos | % of CD8pos | |
| Effector memory CD8 T-cells | CD45RAneg CD197neg CD8pos | % of CD8pos | |
| Terminal differentiated CD8 T-cells | CD57pos CD8pos | % of CD8pos | |
| Exhausted terminal differentiated CD8 T-cells | CD57pos CD279pos CD8pos | % of CD8pos | |
| **Tube 4: TCRgd/TCRab/HLA-DR/CD366/TCR Vd1/CD8b/CD4/CD3/TCR Vd2/CD8** | | | |
| HLA-DRpos CD4 T-cells | HLA-DR-MFI CD4pos | % of CD4pos | HLA-DR_MFI |
| CD366pos CD4 T-cells | CD366MFI CD4pos | % of CD4pos | CD366_MFI |
| HLA-DRpos CD8 T-cells | HLA-DR-MFI CD8pos | % of CD8pos | HLA-DR_MFI |
| CD366pos CD8 T-cells | CD366MFI CD8pos | % of CD8pos | CD366_MFI |
| **TCRγδ T-cells** | TCRγδ-pos CD3pos | % of CD3poa | |
| TCRVδ1 T-cells | TCRVδ1pos TCRγδpos | % of TCRγδ T-cells | |
| TCRVδ2 T-cells | TCRVδ2pos TCRγδpos | % of TCRγδ T-cells | |
| HLA-DRpos monocytes | HLA-DR-MFI CD4dim | % of monocytes | HLA-DR_MFI |
| CD366pos monocytes | CD366MFI CD4dim | % of monocytes | CD366_MFI |
| **Tube 5: Helios/CD127/CD25/CD152/CD39/CD196/FOXP3/CD4/CD161/CD3** | | | |
| Treg | CD25pos CD127neg FOXP3pos HELIOSpos CD4pos | % of CD4pos | |
| CTLA4pos Helios-pos Tregs | CD152MFI FOXP3pos HELIOSpos CD4pos | % of CD4pos | CTLA4_MFI |
| CTLA4pos conventional CD4 T-celler | CD152MFI FOXP3neg HELIOSneg CD4pos | % of CD4pos | CTLA4_MFI |
| Th17 of CD4 T-cells | CD161pos CD196pos CD4pos | % of CD4pos | |
| Tc17 of CD8 T-cells | CD161pos CD196pos CD4neg | % of CD8pos | |
| CTLA4pos CD8 T-cells | CD152MFI CD4neg CD3pos | % of CD8pos | CTLA4_MFI |
| CTLA4pos CD4 T-cells | CD152MFI CD4pos | % of CD4pos | CTLA4_MFI |
| **Tube 6: CD141/LIN/CD304/CD1c/CRLF2/CD301/CD11c/CD123/HLA-DR/CD16** | | | |
| Plasmacytoid dendritic cells (pDC) | CD123pos CD304pos HLA-DRpos LINneg | % of CD45pos | |
| Myeloid dendritic cellc type 1 (mDC1) | CD141pos CD1c-neg HLA-DRpos LINneg | % of CD45pos | |

| Name | Marker combination | Frequency (%) | MFI |
|---|---|---|---|
| Myeloid dendritic cellc type 2 (mDC2) | CD141neg CD1c-pos HLA-DRpos LINneg | % of CD45pos | |
| TSLP-Rpos pDC | CRLF2MFI CD123pos CD304pos HLA-DRpos LINneg | | TSPL-R_MFI |
| TSLP-Rpos mDC1 | CRLF2MFI CD141pos CD1c-neg HLA-DRpos LINneg | | TSPL-R_MFI |
| TSLP-Rpos mDC2 | CRLF2MFI CD141neg CD1c-pos HLA-DRpos LINneg | | TSPL-R_MFI |
| CD301pos pDC | CD301MFI CD123pos CD304pos HLA-DRpos LINneg | | CD301_MFI |
| CD301pos mDC1 | CD301MFI CD141pos CD1c-neg HLA-DRpos LINneg | | CD301_MFI |
| CD301pos mDC2 | CD301MFI CD141neg CD1c-pos HLA-Drpos LINneg | | CD301_MFI |
| **Tube 7: CD35/CD49d/CD10/Singlec-8/CD64/CD66b/CD16/CD11b/HLA-DR/CD14** | | | |
| Granulocytes | CD66b-pos SShigh | | |
| Neutrophils | CD66b-pos Singlec-8neg CD49d-neg SShigh | % of granulocytes | |
| Mature neutrophils | CD10pos CD16pos CD66b-pos Singlec-8neg SShigh | % of neutrophils | |
| Immature neutrophils | CD10neg CD16pos CD66b-pos Singlec-8neg SShigh | % of neutrophils | |
| Metamyelocytes | CD35neg CD49dpos CD66b-pos | % of neutrophils | |
| Neutrophil subpopulation | CD35pos CD49d-pos neutrophils | % of neutrophils | |
| CD11b-dim immature neutrophils | CD11b-dimMFI CD10n CD66b-pos Singlec-8neg CD49d-neg SShigh | % of neutrophils | CD11b_MFI |
| CD11b-bright immature neutrophils | CD11b-brightMFI CD10n CD66b-pos Singlec-8neg CD49d-neg SShigh | % of neutrophils | CD11b_MFI |
| CD11b on all immature neutrophils | CD11bMFI CD10n CD66b-pos Singlec-8neg CD49d-neg | % of neutrophils | CD11b_MFI |
| CD35dim on immature neutrophils | CD35dimMFI CD10n CD66b-pos Singlec-8neg CD49d-neg SShigh | % of neutrophils | CD35_MFI |
| CD35bright on immature neutrophils | CD35brightMFI CD10n CD66b-pos Singlec-8neg CD49d-neg SShigh | % of neutrophils | CD35_MFI |
| CD35 on all immature neutrophils | CD35MFI CD10n CD66b-pos Singlec-8neg CD49d-neg | % of neutrophils | CD35_MFI |
| CD64dim on immature neutrophils | CD64dimMFI CD10n CD66b-pos Singlec-8neg CD49d-neg SShigh | % of neutrophils | CD64_MFI |
| CD64bright on immature neutrophils | CD64brightMFI CD10n CD66b-pos Singlec-8neg CD49d-neg SShigh | % of neutrophils | CD64_MFI |
| CD64 on all immature neutrophils | CD64MFI CD10n CD66b-pos Singlec-8neg CD49d-neg | % of neutrophils | CD64_MFI |
| HLA-DRdim on immature neutrophils | HLA-DRdimMFI CD10n CD66b-pos Singlec-8neg CD49d-neg SShigh | % of neutrophils | HLA-DR_MFI |
| HLA-DRbright on immature neutrophils | HLA-DRbrightMFI CD10n CD66b-pos Singlec-8neg CD49d-neg SShigh | % of neutrophils | HLA-DR_MFI |
| HLA-DR on all immature neutrophils | HLA-DR-MFI CD10n CD66b-pos Singlec-8neg CD49d-neg SShigh | % of neutrophils | HLA-DR_MFI |
| CD49d on all immature neutrophils | CD49dMFI CD10n CD66b-pos Singlec-8neg CD49d-neg | % of neutrophils | CD49d_MFI |
| CD16-dim on immature neutrophils | CD16dimMFI CD10n CD66b-pos Singlec-8neg CD49d-neg | % of neutrophils | CD16_MFI |
| CD16-bright on immature neutrophils | CD16brightMFI CD10n CD66b-pos Singlec-8neg CD49d-neg | % of neutrophils | CD16_MFI |
| CD16 on all immature neutrophils | CD16MFI CD10n CD66b-pos Singlec-8neg CD49d-neg | % of neutrophils | CD16_MFI |
| CD11b-dim on mature neutrophils | CD11b-dimMFI CD10p CD66b-pos Singlec-8neg CD49d-neg SShigh | % of neutrophils | CD11b_MFI |
| CD11b-bright on mature neutrophils | CD11b-brightMFI CD10p CD66b-pos Singlec-8neg CD49d-neg SShigh | % of neutrophils | CD11b_MFI |
| CD11b on all mature neutrophils | CD11bMFI CD10p CD66b-pos Singlec-8neg CD49d-neg | % of neutrophils | CD11b_MFI |
| CD35dim on mature neutrophils | CD35dimMFI CD10p CD66b-pos Singlec-8neg CD49d-neg SShigh | % of neutrophils | CD35_MFI |

| Name | Marker combination | Frequency (%) | MFI |
|---|---|---|---|
| CD35bright on mature neutrophils | CD35brightMFI CD10p CD66b-pos Singlec-8neg CD49d-neg SShigh | % of neutrophils | CD35_MFI |
| CD35 on all mature neutrophils | CD35MFI CD10p CD66b-pos Singlec-8neg CD49d-neg | % of neutrophils | CD35_MFI |
| CD64dim on mature neutrophils | CD64dimMFI CD10p CD66b-pos Singlec-8neg CD49d-neg SShigh | % of neutrophils | CD64_MFI |
| CD64bright on mature neutrophils | CD64brightMFI CD10p CD66b-pos Singlec-8neg CD49d-neg SShigh | % of neutrophils | CD64_MFI |
| CD64 on all mature neutrophils | CD64MFI CD10p CD66b-pos Singlec-8neg CD49d-neg | % of neutrophils | CD64_MFI |
| HLA-DRdim on mature neutrophils | HLA-DRdimMFI CD10p CD66b-pos Singlec-8neg CD49d-neg SShigh | % of neutrophils | HLA-DR_MFI |
| HLA-DRbright on mature neutrophils | HLA-DRbrightMFI CD10p CD66b-pos Singlec-8neg CD49d-neg SShigh | % of neutrophils | HLA-DR_MFI |
| HLA-DR on all mature neutrophils | HLA-DR-MFI CD10p CD66b-pos Singlec-8neg CD49d-neg SShigh | % of neutrophils | HLA-DR_MFI |
| CD16-dim on mature neutrophils | CD16dimMFI CD10p CD66b-pos Singlec-8neg CD49d-neg | % of neutrophils | CD16_MFI |
| CD16-bright on mature neutrophils | CD16brightMFI CD10p CD66b-pos Singlec-8neg CD49d-neg | % of neutrophils | CD16_MFI |
| CD16 on all mature neutrophils | CD16MFI CD10p CD66b-pos Singlec-8neg CD49d-neg | % of neutrophils | CD16_MFI |
| Eosinophils | CD49d-pos Siglec-8pos CD66b-pos SShigh | % of granulocytes | |
| Monocytes | CD14pos | | |
| Monocytes classical | CD14pos CD16neg | % of monocytes | |
| Monocytes intermediate | CD14pos CD16pos | % of monocytes | |
| Monocytes non-classical | CD14neg CD16pos | % of monocytes | |
| CD11b-dim on Monocytes | CD11b-dimMFI CD14pos | % of monocytes | CD11b_MFI |
| CD11b-bright on Monocytes | CD11b-brightMFI CD14pos | % of monocytes | CD11b_MFI |
| CD35dim on Monocytes | CD35dimMFI CD14pos | % of monocytes | CD35_MFI |
| CD35bright on Monocytes | CD35brightMFI CD14pos | % of monocytes | CD35_MFI |
| CD49d-dim on Monocytes | CD49d-dimMFI CD14pos | % of monocytes | CD49d_MFI |
| CD49d-bright on Monocytes | CD49d-brightMFI CD14pos | % of monocytes | CD49d_MFI |
| CD64dim on Monocytes | CD64dimMFI CD14pos | % of monocytes | CD64_MFI |
| CD64bright on Monocytes | CD64brightMFI CD14pos | % of monocytes | CD64_MFI |
| HLA-DRdim on Monocytes | HLA-DRdimMFI CD14pos | % of monocytes | HLA-DR_MFI |
| HLA-DRbright on Monocytes | HLA-DRbrightMFI CD14pos | % of monocytes | HLA-DR_MFI |
| HLA-DR on all Monocytes | HLA-DR-MFI CD14pos | % of monocytes | HLA-DR_MFI |
| CD11b-dim on classical Monocytes | CD11b-dimMFI CD14pos CD16neg | % of classical monocytes | CD11b_MFI |
| CD11b-bright on classical Monocytes | CD11b-brightMFI CD14pos CD16neg | % of classical monocytes | CD11b_MFI |
| CD11b on all classical Monocytes | CD11bMFI CD14pos CD16neg | % of classical monocytes | CD11b_MFI |
| CD35dim on classical Monocytes | CD35dimMFI CD14pos CD16neg | % of classical monocytes | CD35_MFI |
| CD35bright on classical Monocytes | CD35brightMFI CD14pos CD16neg | % of classical monocytes | CD35_MFI |
| CD35 on all classical Monocytes | CD35MFI CD14pos CD16neg | % of classical monocytes | CD35_MFI |
| CD49d-dim on classical Monocytes | CD49d-dimMFI CD14pos CD16neg | % of classical monocytes | CD49d_MFI |
| CD49d-bright on classical Monocytes | CD49d-brightMFI CD14pos CD16neg | % of classical monocytes | CD49d_MFI |
| CD49d on all classical Monocytes | CD49MFI CD14pos CD16neg | % of classical monocytes | CD49d_MFI |

| Name | Marker combination | Frequency (%) | MFI |
|---|---|---|---|
| CD64dim on classical Monocytes | CD64dimMFI CD14pos CD16neg | % of classical monocytes | CD64_MFI |
| CD64bright on classical Monocytes | CD64brightMFI CD14pos CD16neg | % of classical monocytes | CD64_MFI |
| CD64 on all classical Monocytes | CD64MFI CD14pos CD16neg | % of classical monocytes | CD64_MFI |
| HLA-DRdim on classical Monocytes | HLA-DRdimMFI CD14pos CD16neg | % of classical monocytes | HLA-DR_MFI |
| HLA-DRbright on classical Monocytes | HLA-DRbrightMFI CD14pos CD16neg | % of classical monocytes | HLA-DR_MFI |
| HLA-DR on all classical Monocytes | HLA-DR-MFI CD14pos CD16neg | % of classical monocytes | HLA-DR_MFI |
| CD16-dim on classical Monocytes | CD16dimMFI CD14pos CD16neg | % of classical monocytes | CD16_MFI |
| CD16-bright on classical Monocytes | CD16brightMFI CD14pos CD16neg | % of classical monocytes | CD16_MFI |
| CD16 on all classical Monocytes | CD16MFI CD14pos CD16neg | % of classical monocytes | CD16_MFI |
| CD11b-dim on intermediate Monocytes | CD11b-dimMFI CD14pos CD16pos | % of intermediate monocytes | CD11b_MFI |
| CD11b-bright on intermediate Monocytes | CD11b-brightMFI CD14pos CD16pos | % of intermediate monocytes | CD11b_MFI |
| CD11b on all intermediate Monocytes | CD11bMFI CD14pos CD16pos | % of intermediate monocytes | CD11b_MFI |
| CD35dim on intermediate Monocytes | CD35dimMFI CD14pos CD16pos | % of intermediate monocytes | CD35_MFI |
| CD35bright on intermediate Monocytes | CD35brightMFI CD14pos CD16pos | % of intermediate monocytes | CD35_MFI |
| CD35 on all intermediate Monocytes | CD35MFI CD14pos CD16pos | % of intermediate monocytes | CD35_MFI |
| CD49d-dim on intermediate Monocytes | CD49d-dimMFI CD14pos CD16pos | % of intermediate monocytes | CD49d_MFI |
| CD49d-bright on intermediate Monocytes | CD49dMFI CD14pos CD16pos | % of intermediate monocytes | CD49d_MFI |
| CD49d on all intermediate Monocytes | CD49d-brightMFI CD14pos CD16pos | % of intermediate monocytes | CD49d_MFI |
| CD64dim on intermediate Monocytes | CD64dimMFI CD14pos CD16pos | % of intermediate monocytes | CD64_MFI |
| CD64bright on intermediate Monocytes | CD64brightMFI CD14pos CD16pos | % of intermediate monocytes | CD64_MFI |
| CD64 on all intermediate Monocytes | CD64MFI CD14pos CD16pos | % of intermediate monocytes | CD64_MFI |
| HLA-DRdim on intermediate Monocytes | HLA-DRdimMFI CD14pos CD16pos | % of intermediate monocytes | HLA-DR_MFI |
| HLA-DRbright on intermediate Monocytes | HLA-DRbrightMFI CD14pos CD16pos | % of intermediate monocytes | HLA-DR_MFI |
| HLA-DR on all intermediate Monocytes | HLA-DR-MFI CD14pos CD16pos | % of intermediate monocytes | HLA-DR_MFI |
| CD16-dim on intermediate Monocytes | CD16dimMFI CD14pos CD16pos | % of intermediate monocytes | CD16_MFI |
| CD16-bright on intermediate Monocytes | CD16brightMFI CD14pos CD16pos | % of intermediate monocytes | CD16_MFI |
| CD16 on all intermediate Monocytes | CD16MFI CD14pos CD16pos | % of non-classical monocytes | CD16_MFI |
| CD11b-dim on non-classical Monocytes | CD11b-dimMFI CD14neg CD16pos | % of non-classical monocytes | CD11b_MFI |
| CD11b-bright on non-classical Monocytes | CD11b-brightMFI CD14neg CD16pos | % of non-classical monocytes | CD11b_MFI |
| CD11b on all non-classical Monocytes | CD11bMFI CD14neg CD16pos | % of non-classical monocytes | CD11b_MFI |
| CD35dim on non-classical Monocytes | CD35dimMFI CD14neg CD16pos | % of non-classical monocytes | CD35_MFI |
| CD35bright on non-classical Monocytes | CD35brightMFI CD14neg CD16pos | % of non-classical monocytes | CD35_MFI |
| CD35 on all non-classical Monocytes | CD35MFI CD14neg CD16pos | % of non-classical monocytes | CD35_MFI |
| CD49d-dim on non-classical Monocytes | CD49d-dimMFI CD14neg CD16pos | % of non-classical monocytes | CD49d_MFI |
| CD49d-bright on non-classical Monocytes | CD49d-brightMFI CD14neg CD16pos | % of non-classical monocytes | CD49d_MFI |

| Name | Marker combination | Frequency (%) | MFI |
|---|---|---|---|
| CD49d on all non-classical Monocytes | CD49dMFI CD14neg CD16pos | % of non-classical monocytes | CD49d_MFI |
| CD64dim on non-classical Monocytes | CD64dimMFI CD14neg CD16pos | % of non-classical monocytes | CD64_MFI |
| CD64bright on non-classical Monocytes | CD64brightMFI CD14neg CD16pos | % of non-classical monocytes | CD64_MFI |
| CD64 on all non-classical Monocytes | CD64MFI CD14neg CD16pos | % of non-classical monocytes | CD64_MFI |
| HLA-DRdim on non-classical Monocytes | HLA-DRdimMFI CD14neg CD16pos | % of non-classical monocytes | HLA-DR_MFI |
| HLA-DRbright on non-classical Monocytes | HLA-DRbrightMFI CD14neg CD16pos | % of non-classical monocytes | HLA-DR_MFI |
| HLA-DR on all non-classical Monocytes | HLA-DR-MFI CD14neg CD16pos | % of non-classical monocytes | HLA-DR_MFI |
| CD16-dim on non-classical Monocytes | CD16dimMFI CD14neg CD16pos | % of non-classical monocytes | CD16_MFI |
| CD16-bright on non-classical Monocytes | CD16brightMFI CD14neg CD16pos | % of non-classical monocytes | CD16_MFI |
| CD16 on all non-classical Monocytes | CD16MFI CD14neg CD16pos | % of non-classical monocytes | CD16_MFI |
| mMDSC | HLA-DRlow CD11bpos CD16low CD14pos CD66bneg | % of granulocytes | |
| pmnMDSC | HLA-DRlow CD11bpos CD16low CD14neg CD66bpos | % of granulocytes | |

**Supplementary Table 4. Comparison of cystic fibrosis patients' demographics based on clusters defined by immune cell absolute concentrations.**

| Demographics | Cluster 1 | Cluster 2 | *P* value | Test |
|---|---|---|---|---|
| **All** | 15 | 13 | | |
| **Sex** | | | | |
| Female, n (%) | 8 (53.3) | 3 (23.1) | 0.137 | Fisher's exact |
| Male, n (%) | 7 (46.7) | 10 (76.9) | | |
| **Genotype** | | | | |
| ΔF508 heterozygous, n (%) | 9 (60) | 10 (76.9) | 0.492 | Chi Square |
| ΔF508 homozygous, n (%) | 5 (33.3) | 3 (23.1) | | |
| Other, n (%) | 1 (6.7) | 0 | | |
| **CF-related diabetes** | | | | |
| Yes, n (%) | 3 (20) | 6 (46.2) | 0.228 | Fisher's exact |
| No, n (%) | 12 (80) | 7 (53.8) | | |
| **Age, median (range)** | 24 (12-62) | 44 (17-57) | 0.053 | Mann-Whitney U |
| **FEV1%, median (range)** | 73.43 (46-100) | 41 (21-93.3) | 0.005 | Mann-Whitney U |
| **Total IgG, median (range)** | 13.1 (5.3-17.9) | 13.3 (7.9-24-10) | 0.433 | Mann-Whitney U |
| **BMI, median (range)** | 20.6 (17-24.6) | 20.3 (19.1-24.8) | 0.356 | Mann-Whitney U |
| **Years from first PA isolate, median (range)** | 6.8 (0-49.9) | 30 (0-47.3) | 0.0744 | Mann-Whitney U |

PA = *Pseudomonas aeruginosa*
